# Depot-Specific Roles for C/EBPα in White Adipose Tissue Development and Metabolism

**DOI:** 10.1101/2025.07.22.666007

**Authors:** Krista Y. Hu, Esme A. Dodge, Olivia A. B. Maguire, Katherine Y. Ma, Caio V. Matias, Hector S. Himede, Juliana Gomez Pardo, Miriam Cepeda, Robert C. Bauer

**Affiliations:** Cardiometabolic Genomics Program, Division of Cardiology, Department of Medicine, Columbia University, New York, NY 10032

## Abstract

Rates of obesity and its associated metabolic comorbidities continue to rise in the developed world. It is well established that in obesity, the distribution and not just amount of excess white adipose tissue (WAT) correlates with a person’s risk for comorbidities such as coronary artery disease and type 2 diabetes. Thus, understanding the specific mechanisms that drive WAT development in specific adipose depots could elucidate novel mechanisms of metabolic disease. SNPs near the gene *CEBPA* have been associated with multiple cardiometabolic traits by human genome-wide association studies, including waist-to-hip ratio, suggesting that *CEBPA* regulates WAT distribution. *CEBPA* encodes a well characterized transcription factor (C/EBPα) that is long recognized as a master regulator of adipocyte differentiation, yet depot-specific roles for C/EBPα have not been previously described. To further investigate this genetic link, we generated mice with adipocyte-specific *Cebpa* knockout (Cebpa_ASKO) and found that these mice are almost entirely lacking gonadal WAT (gWAT) despite the inguinal WAT (iWAT) being present in near normal amounts. Despite developing, Cebpa_ASKO iWAT contains fewer and larger adipocytes, and fails to expand when challenged with high fat diet. RNA-seq and functional studies demonstrate evidence of altered lipid metabolism and adipocyte function in Cebpa_ASKO iWAT. Finally, Cebpa_ASKO mice have multiple other metabolic phenotypes, including lipid-laden BAT, increased hepatic triglycerides, and increased plasma cholesterol, all of which worsen with prolonged high fat diet feeding. Taken together, these data highlight depot-specific roles for C/EBPα in adipose tissue development, as well as the importance of adipocyte C/EBPα in maintaining metabolic homeostasis.

## Introduction

Rates of obesity, defined as excess accumulation of white adipose tissue (WAT), continue to rise in the developed world [1,2]. Obesity is well known to associate with multiple metabolic comorbidities, including increased risk for cardiovascular disease (CVD), type 2 diabetes (T2D), and metabolic dysfunction-associated steatotic liver disease (MASLD), among others [1]. Interestingly, the distribution of this excess WAT can result in different risks for metabolic disease [3,4]. There are two main types of WAT: subcutaneous white adipose tissue (scWAT) located under the skin, and visceral white adipose tissue (vWAT) located inside the peritoneum in the abdominal area [5]. Humans with excess vWAT have nearly ten times greater risk of poor metabolic outcomes such as CVD and insulin resistance [3,6,7]. Conversely, excess accumulation of scWAT in humans is sometimes referred to as “metabolically healthy obesity,” as it is less associated with poor metabolic outcomes [3,8]. Given these observations, identifying mechanisms that regulate the distribution of adipose tissue between scWAT and vWAT may elucidate factors that can be leveraged in the treatment of obesity and cardiometabolic disease.

Genome-wide association studies (GWAS) have identified more than 1000 loci that associate with obesity traits in humans, including associations with body mass index (BMI), body fat percentage, waist-to-hip-ratio (WHR), and WHR adjusted for BMI (WHRadjBMI), the last of which is a measure of visceral adiposity [9]. The 19q13 GWAS locus contains SNPs that are associated with numerous cardiometabolic traits including plasma triglycerides, plasma total cholesterol, T2D, BMI, and WHRadjBMI [10,11]. There are multiple genes in the 19q13 GWAS locus, including CCAAT/enhancer binding protein alpha (*CEBPA*) (Supplementary Figure 1) [10]. *CEBPA* encodes for the protein C/EBPα, which is a transcription factor that, in coordination with C/EBPβ and peroxisome proliferator-activated receptor gamma (PPARγ), governs the adipocyte differentiation process. In particular, C/EBPα is known to work in a positive regulatory loop with PPARγ to drive the adipogenic gene expression program [12–14]. In mice, germline whole body *Cebpa* knockout is perinatal lethal, but restoration of hepatic *Cebpa* expression permits survival and the resulting mice are lipodystrophic and lack WAT [15]. While clearly required for adipogenesis, it is unclear how C/EBPα regulates adipose distribution and visceral adiposity specifically.

Given the GWAS association with WHRadjBMI, we hypothesized that adipocyte *Cebpa* plays a role in WAT distribution. To test this hypothesis, we crossed Cebpa_fl/fl mice with AdipoQ-Cre mice, establishing mice with <u>a</u>dipocyte-<u>s</u>pecific <u>k</u>nock<u>o</u>ut of *Cebpa* (Cebpa_ASKO). Here we report that Cebpa_ASKO mice have a near complete absence of gonadal WAT (gWAT), the largest vWAT depot in mice, due to a failure in gWAT development. In stark contrast, Cebpa_ASKO iWAT is present in relatively normal quantities but has drastically altered transcriptome and function. Lean Cebpa_ASKO mice also have multiple other metabolic phenotypes including reduced lipolysis and plasma adiponectin, an inability to expand adipose depots when challenged with high fat diet (HFD), increased brown adipose tissue (BAT) mass and lipid content, and increased hepatic triglycerides and plasma cholesterol. Taken together, these data illuminate a novel adipose depot-specific role C/EBPα and highlight its central role in whole body metabolism.

## Results

### Cebpa_ASKO mice have reduced gWAT mass due to halted development

We generated Cebpa_ASKO mice by crossing Cebpa floxed mice (Cebpa_f/fl) with transgenic mice expressing Cre recombinase under control of the *Adipoq* promoter. As *Adipoq* is not expressed until late in the adipocyte differentiation process [13], Cebpa_ASKO mice represent a model of *Cebpa* knockout during late adipocyte differentiation (Figure 1A). We confirmed efficient knockout of *Cebpa* expression in 10-week-old Cebpa_ASKO gWAT (Figure 1B). Interestingly, 10wk-old Cebpa_ASKO mice do not display any outward phenotypes and have no significant difference in body weight compared to Cebpa_fl/fl controls (Figure 1C). However, upon sacrifice and tissue dissection, we immediately noticed a large reduction in gWAT mass in Cebpa_ASKO mice (Figure 1D), despite the normal appearance of other WAT depots (Supplementary Figure 2). This finding was true in both male and female Cebpa_ASKO mice (Figure 1E-F, Supplementary Figure 3). Together, these data clearly demonstrate a gWAT-specific role for C/EBPα in WAT development.

**Figure 1.**
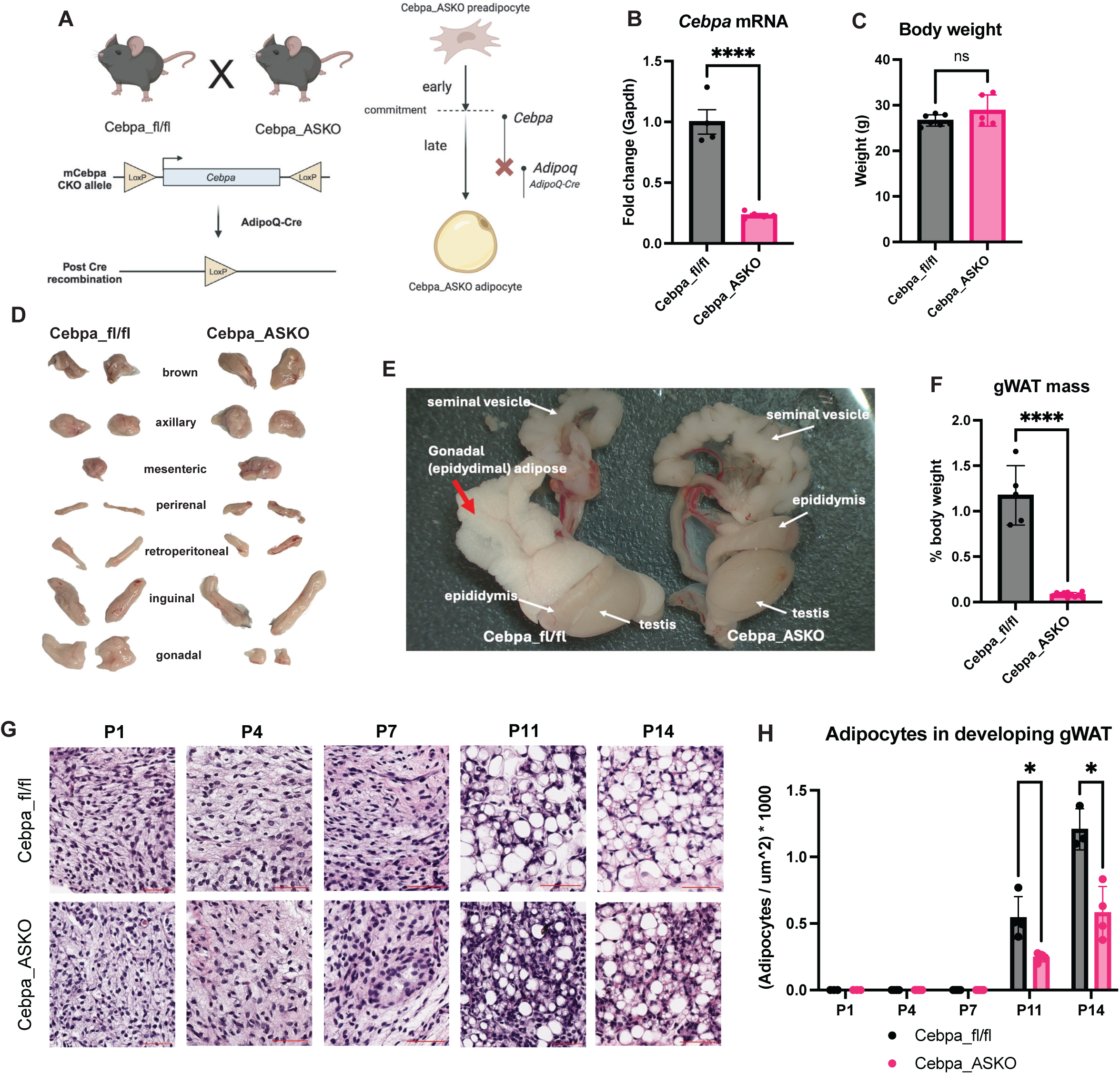
Cebpa_ASKO mice have altered gWAT development. **A.** Schematic of Cebpa_ASKO mouse alleles and adipocyte differentiation KO timeline. **B.** Taqman qPCR for Cebpa in gWAT from male 10-week-old Cebpa_fl/fl and Cebpa_ASKO mice (n=4-5). **C.** Body weight in male 10-12-week-old Cebpa_fl/fl and Cebpa_ASKO mice (n=5-7). **D.** Macroscopic image of adipose tissue depots from male 10-week-old Cebpa_fl/fl and Cebpa_ASKO mice. **E.** Macroscopic image of gWAT attached to gonads from male 10-week-old Cebpa_fl/fl and Cebpa_ASKO mice. **F.** Mass of gWAT from male 10-12-week-old Cebpa_fl/fl and Cebpa_ASKO mice (n=5-7). **G.** H&E stain of developing gWAT appendage from Cebpa_fl/fl and Cebpa_ASKO neonates. **H.** Identification and quantification of adipocytes by Adiposoft in one section of developing gWAT appendages. All mice were chow-fed. Student’s t-test was used to analyze results (*p<0.05, **p<0.01, ***p<0.001, ****p<0.0001). Schematic created with Biorender.

gWAT is a visceral WAT depot that develops postnatally [16], unlike the subcutaneous iWAT that develops in utero. A previous study showed that male mouse gWAT develops during the first two postnatal weeks from an appendage underneath the corpus epididymis, with adipocytes first appearing around P7 [17]. Therefore, we hypothesized that Cebpa_ASKO gWAT development from the appendage diverges from Cebpa_fl/fl mice during this time. To test this, we performed a time-course dissection of the developing appendage at postnatal day 1 (P1), P4, P7, P11, and P14 in both Cebpa_fl/fl and Cebpa_ASKO male mice. H&E staining in both groups revealed the appearance of adipocytes in the developing appendage between days P7 and P11 (Figure 1G). However, by P14, Cebpa_ASKO gWAT appendages have fewer lipid droplets than Cebpa_fl/fl gWAT appendages, consistent with disrupted adipocyte differentiation (Figure 1H). Thus, the severely decreased gWAT mass seen in 10-week-old Cebpa_ASKO mice is due to its halted development in the second perinatal week.

### Cebpa_ASKO iWAT adipocytes develop despite loss of Cebpa

In contrast to the gWAT, iWAT from adult Cebpa_ASKO mice appears grossly normal upon dissection despite *Cebpa* deletion (Figure 2A-B). Closer examination revealed that Cebpa_ASKO iWAT exhibits a small but significant reduction in mass compared to control mice (Figure 2C). Furthermore, Cebpa_ASKO iWAT adipocytes are larger than controls on average (Figure 2D-E), yet Cebpa_ASKO iWAT fat pads contain fewer adipocytes than control iWAT (Figure 2F), consistent with the smaller mass of the fat pad overall. To confirm that Cebpa_ASKO iWAT adipocytes can develop in a cell autonomous manner, we cultured and differentiated stromal vascular fraction cells (SVF) isolated from control and Cebpa_ASKO iWAT. In SVF from Cebpa_fl/fl mice, *Cebpa* exhibits a nearly 6-fold increase in expression during differentiation as expected. Conversely, *Cebpa* expression briefly increases and then retreats to baseline levels in differentiating SVF from Cebpa_ASKO animals (Figure 2G), consistent with the expected timing of Cre expression. We observed no difference in the capacity of Cebpa_fl/fl and Cebpa_ASKO iWAT SVF to develop lipid droplets over the course of differentiation (Figure 2H-I), consistent with the observed in vivo phenotype. These findings suggest that iWAT adipocytes are still able to accumulate lipid and develop the typical unilocular lipid droplet morphology of white adipocytes despite lacking *Cebpa* expression during late differentiation.

**Figure 2.**
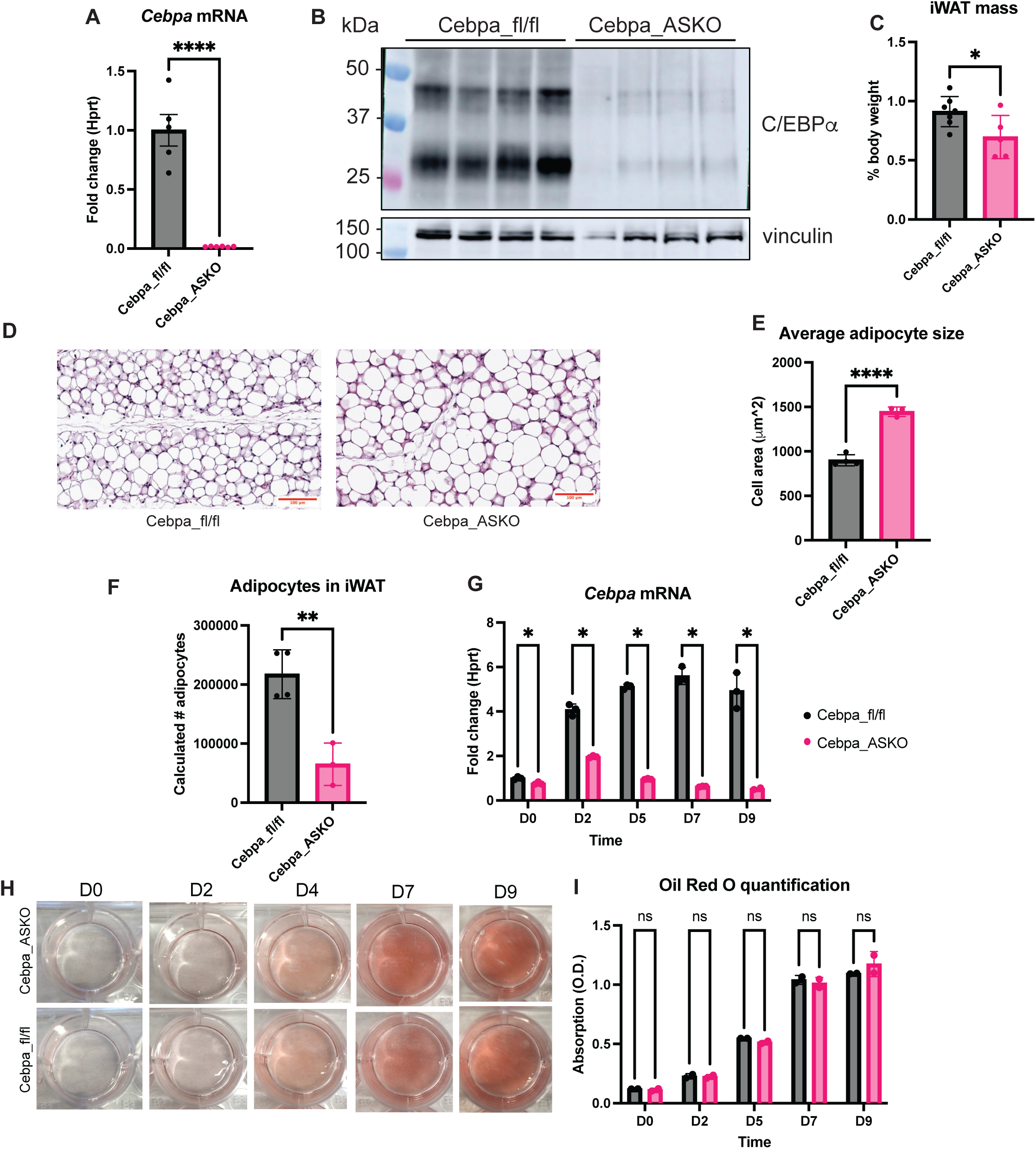
Cebpa_ASKO iWAT develops despite *Cebpa* loss. **A.** Taqman qPCR for *Cebpa* in iWAT from Cebpa_fl/fl and Cebpa_ASKO mice (n=5-7). **B.** Western blot for C/EBPα on iWAT homogenates from 10-week-old male mice. **C.** Mass of iWAT from 10-12-week-old male Cebpa_fl/fl and Cebpa_ASKO mice. **D.** Representative H&E stain of iWAT from 10-week-old male mice. **E, F.** Quantification of cell size by Adiposoft and estimate of adipocyte number by Goldrick formula (n=3-4). **G.** Taqman qPCR for *Cebpa* during differentiation in iWAT SVF. **H.** Neutral lipid droplets during SVF-derived adipocyte differentiation at D0, D2, D5, D7, and D9 visualized by Oil Red O staining. **I.** Quantification of extracted stained lipids from H (n=2). **J.** Leptin measured via ELISA in plasma of 4hr-fasted Cebpa_ASKO mice (n=5-7). Student’s t-test was used to analyze results (*p<0.05, **p<0.01, ****p<0.0001).

### Cebpa_ASKO mice cannot expand adipose depots upon HFD challenge

Given the specific defects in gWAT and iWAT in lean Cebpa_ASKO mice, we next asked if prolonged caloric excess would prompt adipose remodeling and overcome the overall reduction in adipose mass. We challenged Cebpa_fl/fl and Cebpa_ASKO mice with a 60 kcal% high fat diet (HFD) and found that Cebpa_ASKO mice fed HFD for 17 weeks gain significantly less weight than controls (Figure 3A-C). MRI measurements showed that Cebpa_ASKO mice had less fat mass and more lean mass than controls at 15 weeks of HFD feeding (Figure 3D), suggesting that the inability of Cebpa_ASKO mice to gain weight by HFD challenge is due to disrupted adipose expansion. Consistent with this, Cebpa_ASKO mice exhibited a complete absence of adipose depot expansion after 20 weeks on HFD, in stark contrast to the normal expansion of WAT depots in Cebpa_fl/fl mice (Figure 3E, H). Specifically, Cebpa_ASKO gWAT remained nearly completed absent, and Cebpa_ASKO iWAT did not expand (Figure 3F-G). These data suggest that all WAT depots require *Cebpa* expression in mature adipocytes for expansion via hyperplasia or hypertrophy upon HFD challenge.

**Figure 3.**
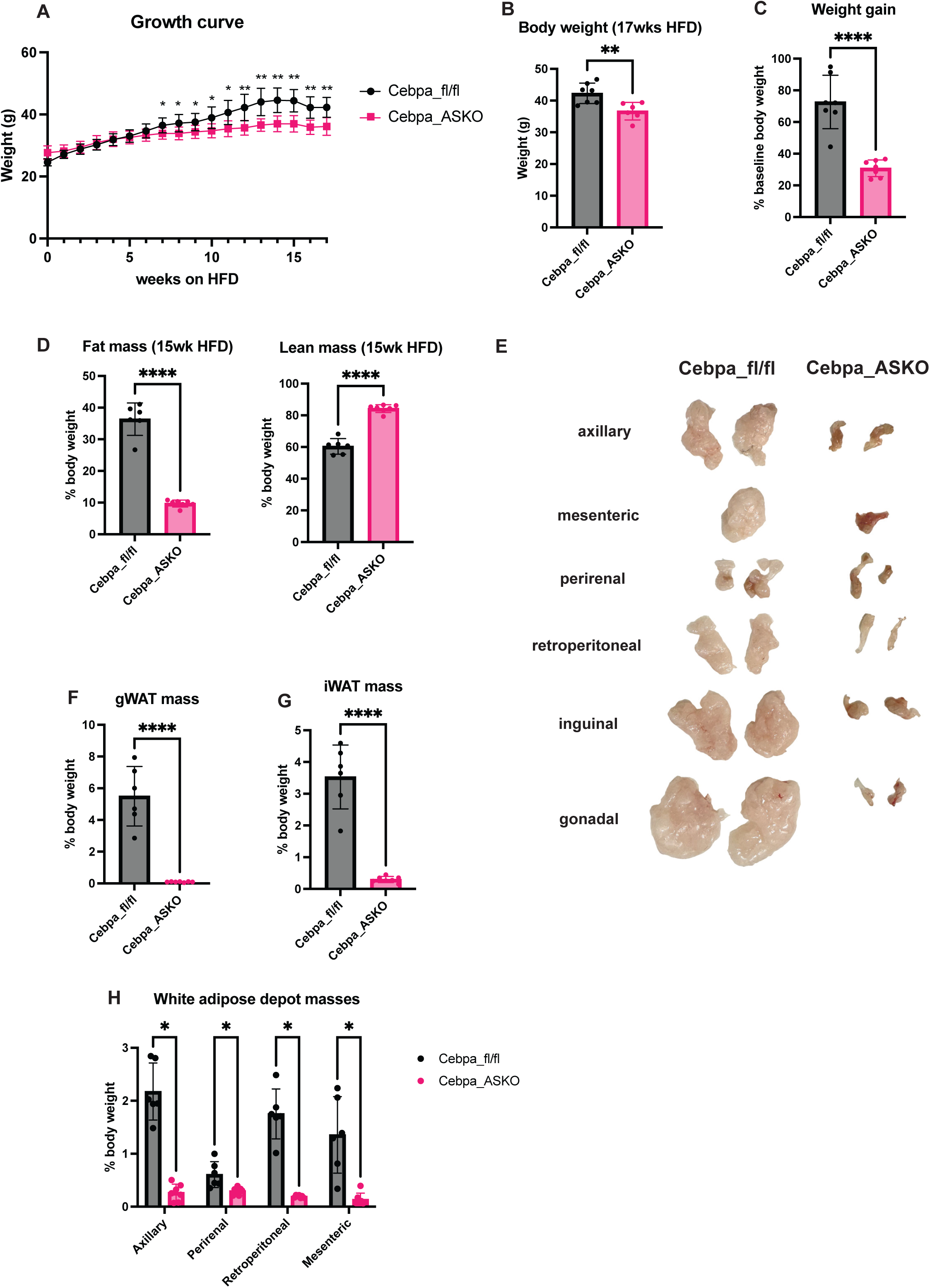
Cebpa_ASKO mice do not expand adipose on a high fat diet. **A.** Growth curve of Cebpa_fl/fl and Cebpa_ASKO male mice fed a high fat diet (HFD) starting at 10-12 weeks of age. **B.** Body weights of Cebpa_fl/fl and Cebpa_ASKO mice after 17 weeks of HFD feeding. **C.** Weight gain in Cebpa_fl/fl and Cebpa_ASKO mice after 17 weeks of HFD feeding measured as percent change in body weight. **D.** Magnetic resonance imaging measured fat mass and lean mass in a separate cohort of Cebpa_fl/fl and Cebpa_ASKO mice after 15 weeks of HFD feeding. **E.** Macroscopic image of adipose depots of Cebpa_fl/fl and Cebpa_ASKO mice after 20 weeks of HFD feeding. **F-H.** Masses of gWAT, iWAT, and other adipose depots from E reported by percent body weight. N=6-7. Student’s t-test was used to analyze results (*p<0.05, ****p<0.0001).

### Cebpa_ASKO iWAT adipocytes have altered transcriptomes and function

The observation that Cebpa_ASKO iWAT depots could not expand after prolonged HFD challenge suggests adipocyte dysfunction, despite their somewhat normal morphology. To further investigate this, we performed bulk RNA-seq on whole iWAT from lean, 10wk-old Cebpa_fl/fl and Cebpa_ASKO mice (Figure 4A-B). Gene set enrichment analysis (GSEA) revealed changes in multiple lipid metabolism processes including fatty acid metabolism and lipid catabolism, consistent with disruption of the canonical adipocyte function of storing and mobilizing lipids (Figure 4C). Interestingly, although iWAT adipocytes have unilocular white adipocyte morphology and iWAT SVF can differentiate, Cebpa_ASKO iWAT has decreased expression of numerous genes important for adipocyte differentiation (*Pparg*, *Zfp423*, *Fabp4*, *Adipoq*) (Supplementary Figure 5). This is consistent with the known role for C/EBPα in transcriptional regulation of differentiation but suggests that a lower expression of these genes is tolerated during iWAT differentiation. Validating our RNA-seq results, *Adipoq* expression is decreased in Cebpa_ASKO iWAT (Figure 4D and F), and Cebpa_ASKO mice also have significantly lower circulating adiponectin compared to controls (Figure 4E), consistent with prior literature showing that *Adipoq* is a downstream target of C/EBPα along with PPARγ [18–20]. However, considering that adiponectin itself is a commonly used marker for mature adipocytes [21], reduced adiponectin circulation and expression again pointed towards disruption of normal adipocyte function in Cebpa_ASKO animals.

**Figure 4.**
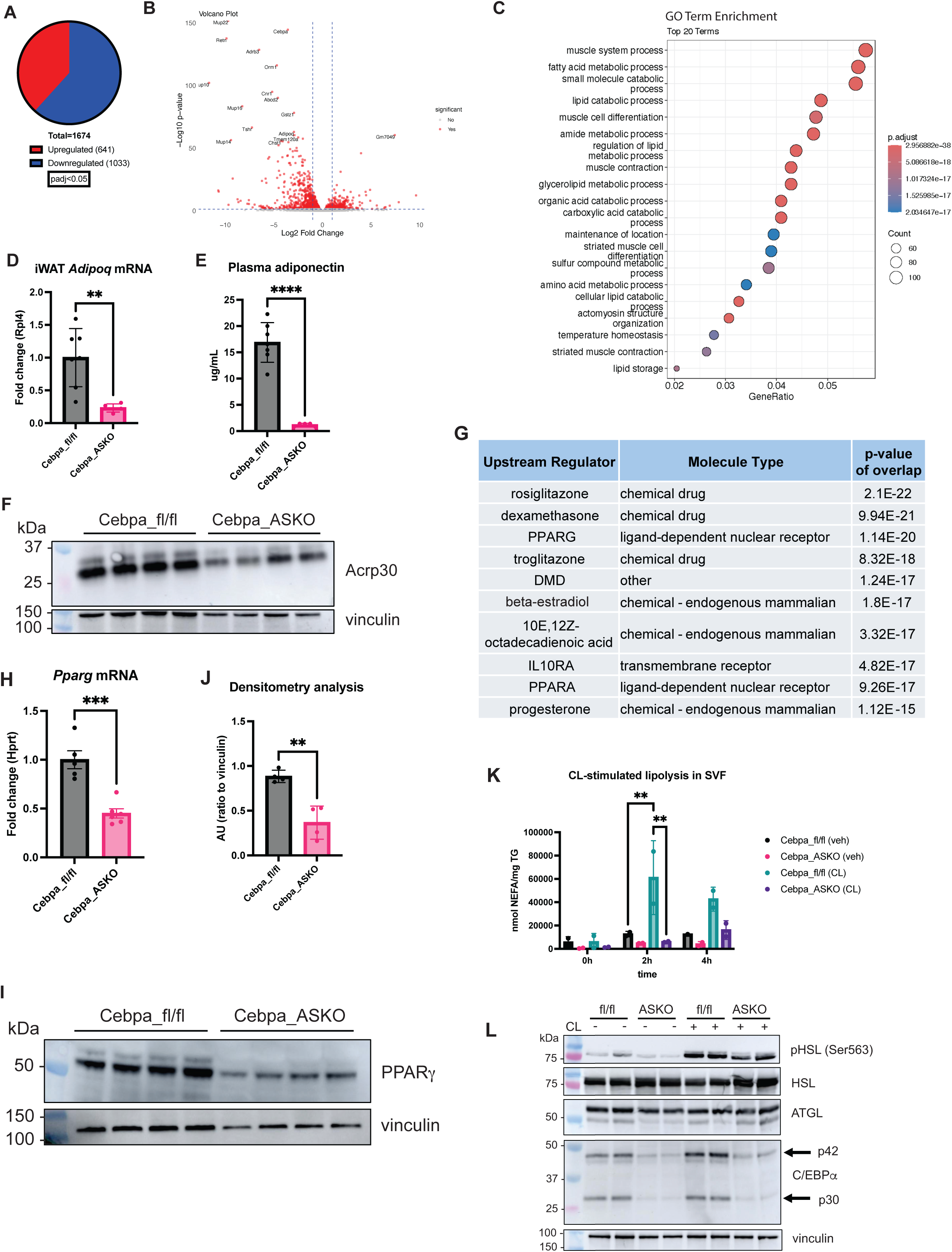
Cebpa_ASKO iWAT is dysfunctional. **A.** Number of differentially expressed genes in bulk RNA-seq of control and Cebpa_ASKO iWAT total tissue RNA. **B.** Volcano plot of DESeq2 analysis of bulk RNA-seq data from Cebpa_fl/fl and Cebpa_ASKO iWAT (n=3). **C.** Gene set enrichment analysis for GO terms from bulk iWAT RNA-seq data. **D.** Taqman qPCR for *Adipoq* in iWAT from Cebpa_fl/fl and Cebpa_ASKO mice (n=5-7). **E.** Adiponectin measured via ELISA in plasma of 4hr-fasted Cebpa_ASKO mice (n=5-7). **F.** Adiponectin (Acrp30) protein levels measured via western blot in lysates from Cebpa_fl/fl and Cebpa_ASKO iWAT. **G.** Identification of upstream regulators most likely to contribute to differences in gene expression using Ingenuity Pathway Analysis. **H.** Taqman qPCR for *Pparg* in whole iWAT (n=5-6). **I.** Western blot analysis of PPARg in homogenates of whole iWAT (n=4). **J.** Densitometry analysis of I, normalized to vinculin. **K.** Non-esterified fatty acid concentration measured in media from iWAT SVF-derived adipocytes treated with vehicle or CL-316,243, normalized to triglyceride mass of each sample (n=2)**. L.** Western blot analysis of lipolysis markers in homogenates of SVF-derived adipocytes from G after 2h of stimulation. All mice were 10-12-weeks-old and chow-fed. Student’s t-test was used to analyze results (*p<0.05, **p<0.01, ****p<0.0001).

Ingenuity Pathway Analysis (IPA) revealed that the most significantly changed upstream regulatory networks in Cebpa_ASKO iWAT were those regulated by PPARγ or PPARγ agonists, suggesting that decreased PPARγ activity is contributing to the reduced function (Figure 4G). *Pparg* expression and protein levels are indeed decreased in whole Cebpa_ASKO iWAT (Figure 4H-J), pointing towards a decrease in PPARγ activity in Cebpa_ASKO iWAT that is consistent with the described positive regulatory loop of C/EBPα and PPARγ [14,22,23]. It is possible that the decrease in PPARγ expression and activity is driving the dysfunction in Cebpa_ASKO adipocytes in conjunction with loss of *Cebpa,* while the residual *Pparg* expression may be sufficient to drive iWAT development in Cebpa_ASKO mice.

Our RNA-seq results also indicated decreased expression of lipolysis genes in Cebpa_ASKO iWAT compared to controls (Supplementary Figure 5), including *Pnpla2*, *Lipe*, and *Mgll*. Lipolysis is a characteristic function of adipocytes [24] and largely recognized to be regulated at the protein level, so decreased transcription of these genes may not result in functional changes [25]. Therefore, we investigated lipolysis in SVF-derived adipocytes from Cebpa_ASKO iWAT. Upon stimulating lipolysis with CL-316,243 in differentiated SVF-derived iWAT adipocytes, we found that release of non-esterified fatty acids (NEFAs) was decreased in Cebpa_ASKO cells compared to Cebpa_fl/fl cells (Figure 4K). Additionally, Western blot analysis showed decreased protein levels of lipolysis markers such as phosphorylated Ser563 in HSL in the Cebpa_ASKO lysates (Figure 4L). These data confirmed that Cebpa_ASKO iWAT adipocytes have decreased lipolysis function compared to Cebpa_fl/fl adipocytes. Taken together, Cebpa_ASKO iWAT demonstrates multiple aspects of dysfunction, including an altered transcriptome, reduced adipokine production, and decreased lipolysis, despite the tissue having a somewhat normal appearance.

### Cebpa_ASKO mice have whiter BAT

Given that the AdipoQ-Cre transgene is expressed in BAT as well, we found that Cebpa_ASKO mice have efficient *Cebpa* knockout in their BAT (Figure 5A) as well. Cebpa_ASKO BAT has significant increased mass (Figure 5B), and H&E staining revealed the Cebpa_ASKO BAT to have excess lipid droplet content (Figure 5C). Consistent with this “whitening,” there is a downward trend in expression of typical brown adipocyte markers (*Ucp1*, *Cidea*, *Ppargc1a*) that does not reach significance. Additionally, expression of leptin, an indicator of whitening and white adipocyte identity [26], was increased in Cebpa_ASKO BAT (Figure 5D), reflecting the appearance of unilocular lipid droplets, a trait of white adipocytes. Mice challenged with 20 weeks of HFD feeding showed progression of increased BAT mass and worsening lipid accumulation (Figure 5E-G). These findings suggest a potential role for C/EBPα in BAT lipid metabolism or storage [15].

**Figure 5.**
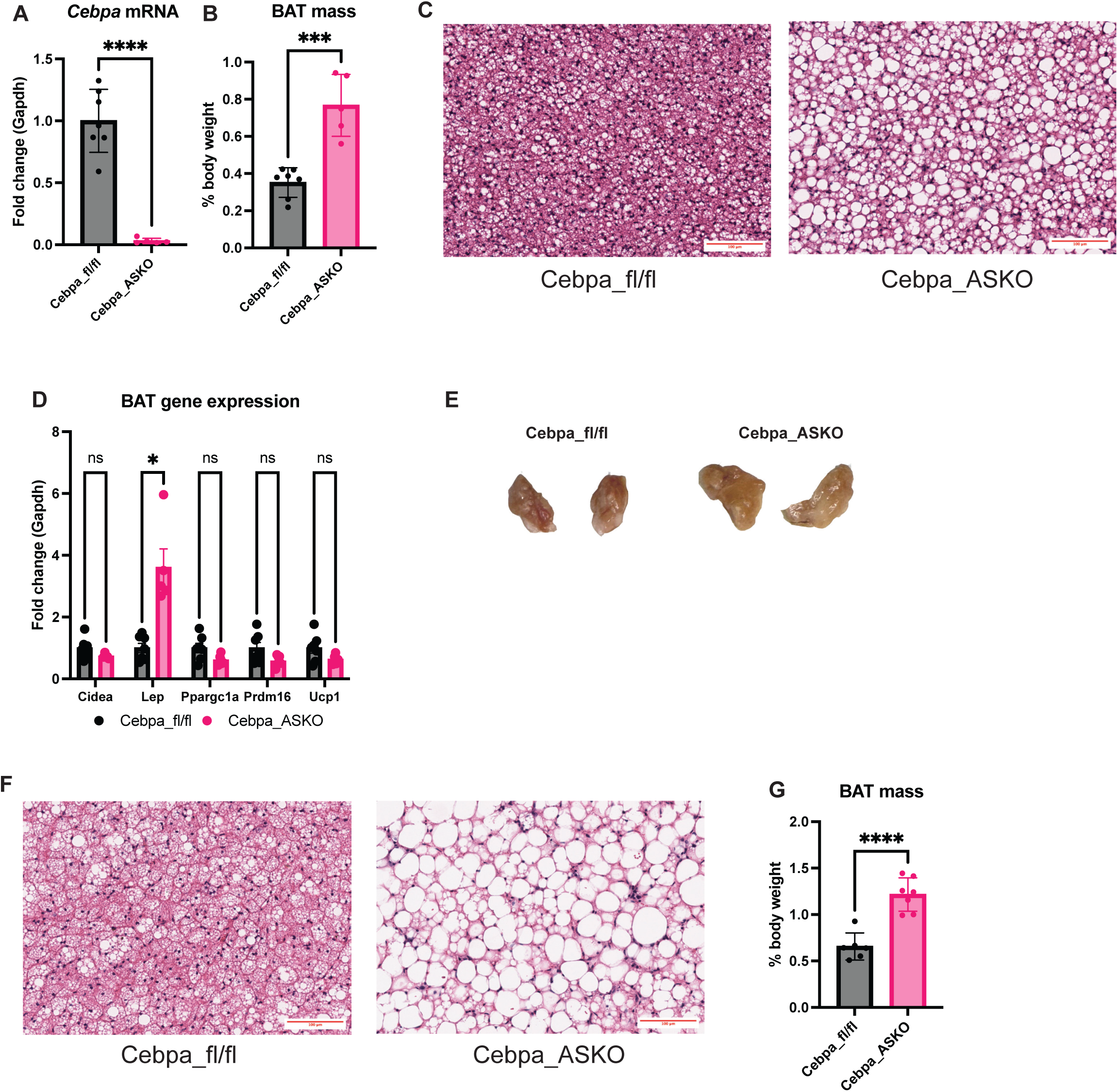
Cebpa_ASKO mice have whitened BAT. **A.** Taqman qPCR for *Cebpa* in RNA extracted from whole BAT (n=5-7). **B.** BAT mass reported as percentage of body weight in Cebpa_fl/fl and Cebpa_ASKO mice. **C.** Representative H&E stain of BAT from Cebpa_fl/fl and Cebpa_ASKO mice. **D.** Taqman qPCR for *Cidea, Lep, Ppargc1a, Prdm16,* and *Ucp1* expression in RNA extracted from whole BAT (n=5-7). **E.** Representative H&E stain of BAT from Cebpa_fl/fl and Cebpa_ASKO mice fed HFD for 20 weeks. **F.** Macroscopic image of BAT from E. **G.** BAT mass reported as percentage of body weight in Cebpa_fl/fl and Cebpa_ASKO mice after 20 weeks of HFD feeding. Male mice were 10-12-weeks-old and chow-fed unless otherwise noted. Student’s t-test was used to analyze results (*p<0.05, ***p<0.001, ****p<0.0001).

### Cebpa_ASKO mice have hypertrophic livers

In both mice and humans, partial and complete lipodystrophies impact hepatic lipid metabolism [27–29]. As our Cebpa_ASKO mice have partial lipodystrophy shown by reduced WAT mass and altered WAT lipid metabolism, we investigated the livers of Cebpa_ASKO mice [30]. Lean Cebpa_ASKO mice have significantly larger livers than Cebpa_fl/fl mice (Figure 6A). They also have increased hepatic triglycerides although hepatic cholesterol remains unchanged (Figure 6B-C). Cebpa_ASKO mice also have increased plasma cholesterol, with no change in plasma triglycerides compared to controls (Figure 6D-E). These phenotypes are all further exacerbated in Cebpa_ASKO mice challenged with HFD feeding (Supplementary Figure 6). Given the increase in plasma cholesterol, we investigated the well-known LDLR/PCSK9 cholesterol metabolic pathway [31]. We found no change in both circulating plasma PCSK9 levels and hepatic protein levels of LDLR (Figure 6F-G). These indicate that the increase in plasma cholesterol in Cebpa_ASKO mice is independent of the PCSK9/LDLR pathway.

**Figure 6.**
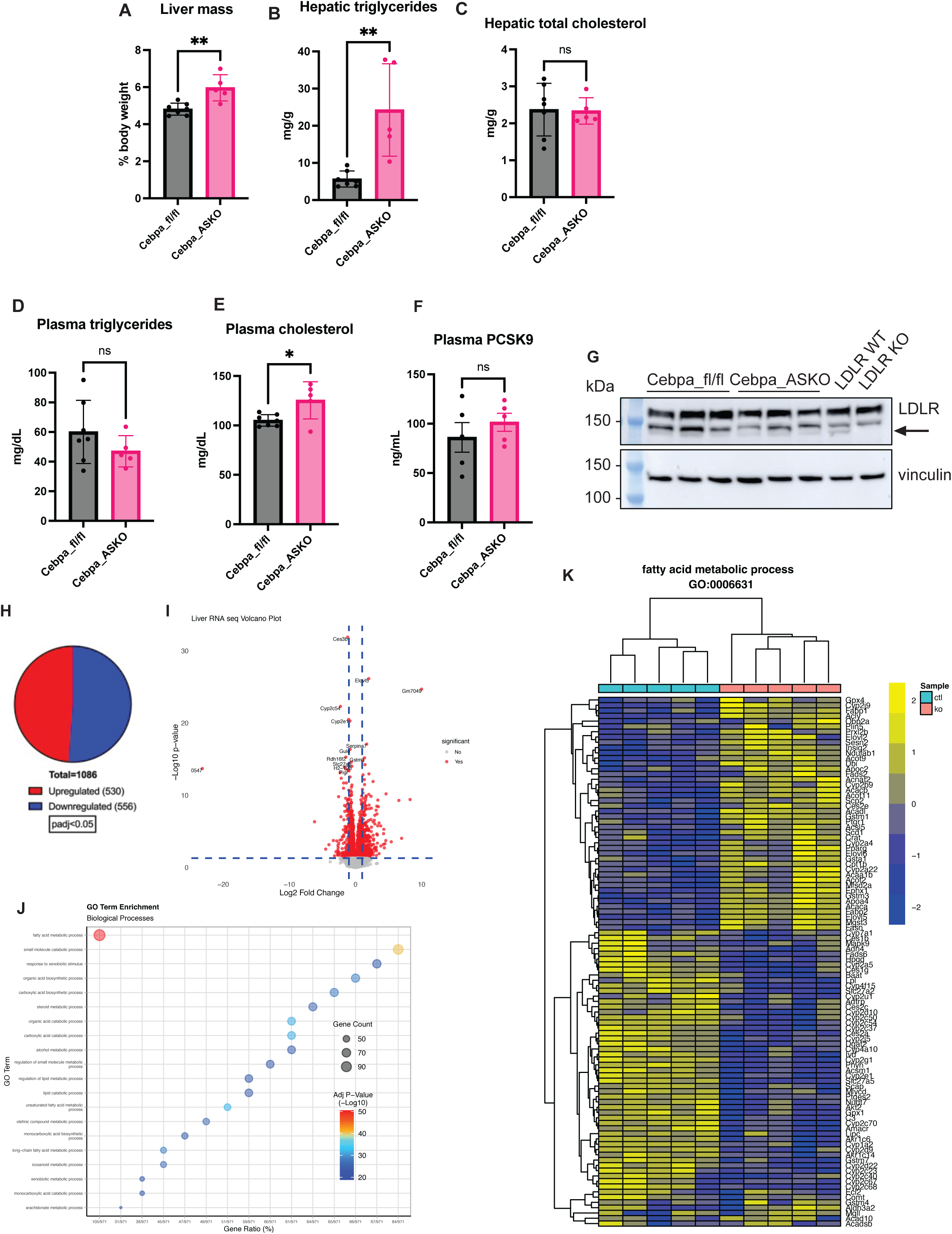
Cebpa_ASKO mice have increased lipid accumulation in their livers. **A.** Liver mass reported as percentage of body weight from Cebpa_fl/fl and Cebpa_ASKO mice. **B.** Hepatic triglycerides and **(C)** cholesterol measured in liver homogenates (n=5-7). **D.** Plasma triglycerides and **(E)** cholesterol measured in 4h fasted mice (n=5-7). **F.** Plasma PCSK9 levels measured via ELISA in 4h fasted mice (n=5). **G.** Western blot analysis of hepatic LDLR protein levels. **H.** Pie chart of number of differentially expressed genes in bulk RNA-seq. **I.** Volcano plot of DESeq2 analysis of bulk RNA-seq data from Cebpa_fl/fl and Cebpa_ASKO livers (n=5). **J.** Gene set enrichment analysis for GO terms from RNA-seq data. **K.** Heatmap of expression of genes clustered in the fatty acid metabolic process GO term from bulk RNA-seq. All mice were 10-12 weeks old and chow-fed. Student’s t-test was used to analyze results (*p<0.05, **p<0.01).

To further investigate the effect on liver lipid metabolism of the multiple adipose depot phenotypes in Cebpa_ASKO mice, we performed bulk RNA-seq on livers from lean, 10-12-week-old Cebpa_fl/fl and Cebpa_ASKO (Figure 6H-I). Gene set enrichment analysis showed fatty acid metabolism to be the most significant gene ontology signature in Cebpa_ASKO livers (Figure 6J), with upregulation of fatty acid metabolism genes such as *Fasn*, *Scd1*, and *Elovl6*, aligning with the increase in hepatic triglycerides (Figure 6K). These findings suggest that increased hepatic TG deposition and synthesis in Cebpa_ASKO livers may be driving altered hepatic lipid and lipoprotein metabolism.

## Discussion

C/EBPα has been well described as a master transcriptional regulator of adipogenesis. Prior studies of mice harboring global germline deletion of *Cebpa* demonstrate a complete lack of WAT development, underscoring the importance of C/EBPα for normal WAT development. Interestingly, GWAS have identified SNPs near *CEBPA* that associate with metrics of visceral adiposity such as waist-to-hip ratio adjusted for BMI (WHRadjBMI), suggesting that C/EBPα plays depot-specific roles in WAT development, yet there is no prior experimental evidence supporting this. Here, we report a novel model of adipocyte-specific *Cebpa* knockout that dramatically reduces gWAT mass, the largest visceral adipose depot in mice, while leaving iWAT somewhat intact, demonstrating a gWAT-specific role for C/EBPα in WAT development. These data are the first in vivo validation of the 19q13 GWAS association with WHRadjBMI and suggest that *CEBPA* is a causal gene at this locus.

The *Cebpa* gene lies in the 19q13 GWAS locus, SNPs in which have been repeatedly associated with metrics of adiposity such as BMI, WHR, WHRadjBMI, visceral adipose tissue volume, and body fat percentage [10,11,32], in addition to other metabolic traits including circulating adiponectin and plasma total cholesterol [33,34]. There are three genes in the 19q13 GWAS locus: *CEBPA*, *PEPD,* and *CEBPG* (Supplemental Figure 1), with most SNPs often annotated to *PEPD* given that some GWAS SNPs lie in the first *PEPD* intron [10,35–42]. A recent report demonstrated that macrophage *Pepd* can regulate adipose fibrosis in mice [43], but experimental evidence supporting *Pepd* or *Cebpg* as the causal gene(s) in this locus is scant, particularly in comparison to the body of literature implicating *Cebpa* in many of the associated phenotypes. Interestingly, many GWAS SNPs in this region demonstrate eQTLs with *PEPD* expression in human adipose tissue per the GTEx consortium, while *CEBPA* is completely devoid of any adipose eQTLs despite its well characterized role in adipose development. Given the multitude of metabolic GWAS associations in this locus, further study is warranted to determine the causal genes and biological mechanisms underlying those associations. Despite the lack of eQTLs, the data presented in this manuscript supports the notion that *CEBPA* is one of the causal genes in the 19q13 GWAS locus.

Although past studies have investigated the effects of acutely induced adipocyte-specific *Cebpa* knockout, none have reported adipose depot-specific effects. Wang et al. reported that inducing *Cebpa* knockout via doxycycline-inducible Cre from embryonic day 11 through postnatal day 16 did not affect gWAT development or mass [44]. Unlike that model, our Cebpa_ASKO mice have knockout of *Cebpa* driven by the constitutively active AdipoQ-Cre transgene, and we suspect that any phenotypic differences between the models must be related to method of knockout. Our use of the AdipoQ-Cre model introduces some complexity in understanding when *Cebpa* knockout occurs, as *Adipoq* is expressed only in the late stages of adipogenesis, and its transcription is driven by C/EBPα, which is active in earlier stages [19,45,46]. Therefore, our interpretation of the Cebpa_ASKO mouse is that *Cebpa* would be expressed in preadipocytes in early differentiation but then silenced near maturation, and we confirmed this with an *in vitro* iWAT SVF time-course experiment (Figure 2 G-I). Since preadipocytes in all Cebpa_ASKO WAT depots presumably express *Cebpa* early in the adipogenic program, the lack of gWAT in Cebpa_ASKO mice suggests that gWAT is more sensitive to loss of C/EBPα in mid to late-stage differentiation than other depots.

The lack of gWAT in Cebpa_ASKO mice aligns with our understanding of C/EBPα as a critical regulator of adipose development, whereas the maintenance of iWAT in those same mice defies expectations. In mice, these depots arise from distinct progenitor cell populations: iWAT develops from the *Prrx1* lineage while gWAT develops from *Wt1*-expressing cells [47–49]. Previous studies have identified a particularly adipogenic cluster of adipocyte progenitor cells (APCs), sometimes termed “committed preadipocytes,” that are present in both adult and developing iWAT and male gWAT [50–52]. Interestingly, this subpopulation expresses *Pparg*, *Adipoq*, *Fabp4* and *Plin1* prior to lipid droplet formation, and they appear fibroblast-like in vitro [50,52–54]. When isolated from embryonic iWAT or perinatal gWAT and cultured in vitro, these cells accumulate lipid and differentiate more efficiently than other populations of progenitor cells [52,54]. A possible explanation for the Cebpa_ASKO phenotype is that there are more “committed preadipocytes” in developing mouse iWAT than gWAT, and these cells are more tolerant to loss of *Cebpa* during differentiation. As we also observe reductions in PPARγ mRNA and protein in Cebpa_ASKO iWAT, this developmental threshold may involve the entire C/EBPα−PPARγ feedback loop. Further studies including identifying changes in APC populations in Cebpa_ASKO developing gWAT will shed light on the mechanisms governing these developmental thresholds. Overall, our findings add to the growing body of literature demonstrating that scWAT and vWAT have different developmental programs and functions despite both being white adipose tissue.

Unlike the two main WAT pads, Cebpa_ASKO BAT is larger and lipid-laden, a phenotype that worsens upon extended HFD feeding. The appearance of large lipid droplets in Cebpa_ASKO BAT is consistent with results from previously reported *Cebpa* knockout models [15,44]. BAT hypertrophy was also seen in the transgenic global *Cebpa* knockout mice [15], confirming that C/EBPα is not necessary for BAT development. Notably, previous data from the inducible adipocyte-specific *Cebpa* knockout model also shows lipid-laden BAT despite no change in WAT mass, suggesting that the whitened appearance of Cebpa_ASKO BAT is not due to the partial lipodystrophy [44]. Therefore, it is possible that C/EBPα acts as a brake on whitening of brown adipocytes during development, leading to whitening upon *Cebpa* loss. The Cebpa_ASKO BAT may also be dysfunctional allowing for lipid accumulation, a possibility reflected in the downward trend of brown adipocyte gene expression. C/EBPα clearly regulates lipid accumulation in BAT, but further investigation is required to understand the exact mechanisms at play.

Finally, Cebpa_ASKO mice also exhibit numerous alterations in hepatic physiology and function, including elevated hepatic triglycerides and plasma cholesterol, and multiple transcriptional changes in hepatic fatty acid metabolism genes. It is plausible that these traits are a result of excess lipid deposition in the liver due to the lack of gWAT in these mice. Furthermore, significant changes in iWAT function, including reduced adipokine secretion and lipolysis, could be affecting liver function via altered crosstalk to the liver, although reduced lipolysis in Cebpa_ASKO mice does not align with increased hepatic TG content. Given ongoing interest in the relationship between vWAT and cardiometabolic traits [7], the mechanisms governing the increased hepatic TG content and plasma cholesterol levels in Cebpa_ASKO mice are the focus of ongoing investigations.

Altogether, these data demonstrate that adipocyte C/EBPα is selectively required for gonadal WAT development in mice. This finding validates the GWAS association of SNPs near *CEBPA* at the 19q13 locus with adiposity metrics such as waist-to-hip-ratio and suggests that C/EBPα contributes to determining adipose distribution in humans. As increased WHR has been well documented as tracking negatively with metabolic traits such as an increased risk for heart disease, understanding the genetic contributions of *CEBPA* and its role in vWAT development brings us one step closer to curbing increasing WHR in the population.

## Materials and Methods

### Animals

Cebpa_fl/fl (Jackson Labs strain #006447) and AdipoQ-Cre + (Jackson Labs strain #010803) mice on the C57BL6/J background were acquired from Jackson Laboratories. These mice were crossed and maintained in-house to generate experimental Cebpa_fl/fl; AdipoQ-Cre + (*Cebpa* adipocyte-specific knockout (Cebpa_ASKO)) and negative control Cebpa_fl/fl; AdipoQ-Cre – (Cebpa_fl/fl) mice. Cebpa_ASKO and Cebpa_fl/fl mice for experiments were bred such that all Cebpa_fl/fl; AdipoQ-Cre + and Cebpa_fl/fl; AdipoQ-Cre – mice were true littermates. Mice were fed a chow diet unless on special diet. Mice were either postnatal days 1-14 (P1-14) or 8-12 weeks old for all experiments unless otherwise indicated. For high fat diet feeding experiments, mice were fed a 60% kcal fat diet from Research Diets (D12492) ad libitum and maintained for 16-20 weeks.

### In vivo experiments

Blood for plasma lipid and ELISA measurements was collected retro-orbitally from anesthetized 4 hour-fasted or random-fed mice with heparinized tubes. All blood samples were then spun at 1500 x g for 10 minutes at 4C to separate plasma. Plasma cholesterol and triglycerides were measured in 4 hour-fasted plasma via plate assay using Infinity Total Cholesterol and Infinity Triglycerides reagents (TR13421 and TR22421). Plasma from random-fed mice or 4h-fasted mice was used for all ELISAs (adiponectin: Millipore-Sigma EZMADP-60K, leptin: Millipore-Sigma EZML-84K, PCSK9: R&D MPC900).

### SVF isolation and culture

Inguinal white adipose tissue (iWAT) was harvested from 3-5 8-10-week-old sex and age-matched mice and placed in ice-cold 1X PBS. Lymph nodes were removed. iWAT was pooled per genotype and minced manually for 5 minutes in digestion media (L-15 Leibovitz media, 1.5% BSA, 1% Pen/Strep, 10 U/mL DNAseI, 480 U/mL hyaluronidase, 0.14 U/mL Liberase TM). Tissue was dissociated by shaking in digestion buffer at 37C for 1 hour at 250 rpm. Homogenate was filtered through a 100 uM cell strainer and spun at 300 x g, 4C, for 10 mins. Supernatant was discarded, and the cell pellet was washed in 10mL cell culture medium (DMEM, 10% FBS, 1% Pen/Strep, 2mM L-Glut), then resuspended in 5mL cell culture medium supplemented with 1 ug/mL insulin. Cell resuspension was plated in a 10cm collagen-coated dish and allowed to grow until 95% confluence, with media replaced every 2-3 days. Then, cells were lifted with 0.25% trypsin, counted, and seeded into either 6-well or 12-well plates. Once cells reached 95% confluence, differentiation was initiated with a cocktail consisting of 10% FBS, 1% Pen/Strep, 5 ug/mL insulin, 1 uM rosiglitazone, 1 uM dexamethasone, and 250 uM IBMX in DMEM/F12. After 48 hours in differentiation cocktail, media was changed to maintenance media consisting of 10% FBS, 1% Pen/Strep, 5 ug/mL insulin, and 1 uM rosiglitazone in DMEM/F12. Experiments were started after day 7 of differentiation unless otherwise noted.

### Oil red O staining

Oil red O (ORO) staining was performed on cell cultures in 12-well plates. Cells were washed twice with 2X PBS before fixing in 10% paraformaldehyde for 30mins at room temperature. Cells were washed twice for with water for 1 min each, then once with 60% isopropanol for 5 mins. Cells were stained with 60% ORO stock solution in water for 15 mins, then washed 3 times with water for 2 mins each before imaging. To quantify ORO area, water was aspirated, and the cells were incubated with 100% isopropanol for 3 mins to dislodge neutral lipids into solution. The isopropanol was then collected and added in equal volumes to a 96-well plate before measuring absorbance at 510nm.

### Bulk RNA-seq of inguinal scWAT and livers

10-week-old female mice were euthanized and 50-80mg of inguinal scWAT was dissected out and placed into ice-cold PBS. Livers were dissected from 10-week-old male mice. Tissues were homogenized in Qiazol Lysis Reagent, and RNA was isolated using the RNeasy Lipid Tissue Mini Kit (Qiagen). Quality was assessed with TapeStation before submission to the Columbia Genome Center for bulk, paired-end RNA-sequencing (NextSeq 500) and differential gene expression analysis (DESeq2).

### Histology

For adipose tissue and liver histology, whole tissues or lobes from Cebpa_fl/fl and Cebpa_ASKO mice and neonates were fixed in 4% PFA for 24 hours before switching to 70% ethanol. Tissues were submitted to the Columbia Molecular Pathology Shared Resource Histology Service, where they were embedded flat in paraffin and sectioned at 7um (neonate tissue) or 8um (adult tissue), for 3 serial sections per slide, followed by H&E staining. The histology service provided whole slide scanning at 40X (Leica SCN 400). Adipose tissue images were analyzed using the Adiposoft ImageJ plugin (parameters: minimum diameter = 10um, maximum diameter = 100um). Adipocyte number calculations were done using the Goldrick formula.

### Molecular assays

#### Quantitative PCR

RNA from adipose and liver tissues was isolated using the RNeasy Lipid Tissue Mini kit (Qiagen). cDNA was synthesized with the High-Capacity cDNA Reverse Transcription Kit (Applied Biosystems). qPCR was performed with pre-designed TaqMan probes from Thermo Fisher Scientific (see Supplementary Information) and TaqMan Fast Advanced Master Mix (Applied Biosystems).

#### Western blot

Cells or tissues were homogenized in RIPA buffer supplemented with 1X Halt Protease and Phosphatase Inhibitor (Thermo Fisher Scientific 78440). Homogenates were clarified by centrifugation at 12,000 x g at 4C for 15mins twice. Protein in clarified homogenates was quantified by BCA assay (Thermo Fisher Scientific), and 20-30ug of protein was run on a premade 10% Bis-Tris SDS-page gel and transferred to a nitrocellulose membrane. Membrane was blocked in 5% milk and incubated in primary antibody overnight (see Supplementary Information). Proteins were detected with HRP-linked secondary antibody and visualized through incubation in Immobilon ECL Ultra Western HRP Substrate (Millipore Sigma WBULS0100) or Luminata Classico Western HRP Substrate (Millipore Sigma WBLUC0500). Membranes were incubated in Restore Western Blot Stripping Buffer (Fisher Scientific PI21059) for 15 min before reblocking and subsequent reprobing.

#### Statistics

Data was graphed using GraphPad Prism 10. Prism 10 was also used to perform 2-tailed Student’s t-tests to compare two groups. Multiple t-tests were used when comparing two groups with multiple variables.

### Supplementary Information

Pre-designed Taqman probes for qPCR

| Probe | Manufacturer | Assay ID |
| --- | --- | --- |
| Taqman *Adipoq* probe | Thermo Fisher Scientific | Mm00456425_m1 |
| Taqman *Cebpa* probe | Thermo Fisher Scientific | Mm00514283_s1 |
| Taqman *Cebpb* probe | Thermo Fisher Scientific | Mm00843434_s1 |
| Taqman *Cidea* probe | Thermo Fisher Scientific | Mm00432554_m1 |
| Taqman *Gapdh* probe | Thermo Fisher Scientific | Mm99999915_g1 |
| Taqman *Hprt* probe | Thermo Fisher Scientific | Mm00446968_m1 |
| Taqman *Lep* probe | Thermo Fisher Scientific | Mm00434759_m1 |
| Taqman *Pparg* probe | Thermo Fisher Scientific | Mm00440940_m1 |
| Taqman *Ppargc1* probe | Thermo Fisher Scientific | Mm01208835_m1 |
| Taqman *Prdm16* probe | Thermo Fisher Scientific | Mm00712556_m1 |
| Taqman *Ucp1* probe | Thermo Fisher Scientific | Mm01244861_m1 |

Antibodies

| Target | Manufacturer | Catalog number | Dilution |
| --- | --- | --- | --- |
| C/EBPα | Cell Signaling Technology | 8178S | 1:1000 |
| ACRP30 | R&D Systems | AF1119 | 1:1000 |
| PPARγ | Cell Signaling Technology | 2443S | 1:1000 |
| Phospho-HSL (Ser563) | Cell Signaling Technology | 4139S | 1:1000 |
| HSL | Cell Signaling Technology | 4107T | 1:1000 |
| ATGL | Cell Signaling Technology | 2138S | 1:1000 |
| Vinculin | Sigma-Aldrich | V9131 | 1:1000 |
| LDLR | Abcam | AB30532 | 1:1000 |
| IgG (H+L) Goat anti-mouse, HRP | Invitrogen | PI31430 | 1:3000 |
| IgG (H+L) Goat anti-rabbit, HRP | Invitrogen | PI31460 | 1:3000 |
| IgG (H+L) Rabbit anti-goat, HRP | Invitrogen | PI31402 | 1:7000 |

---

**Supplementary Figure 1:**
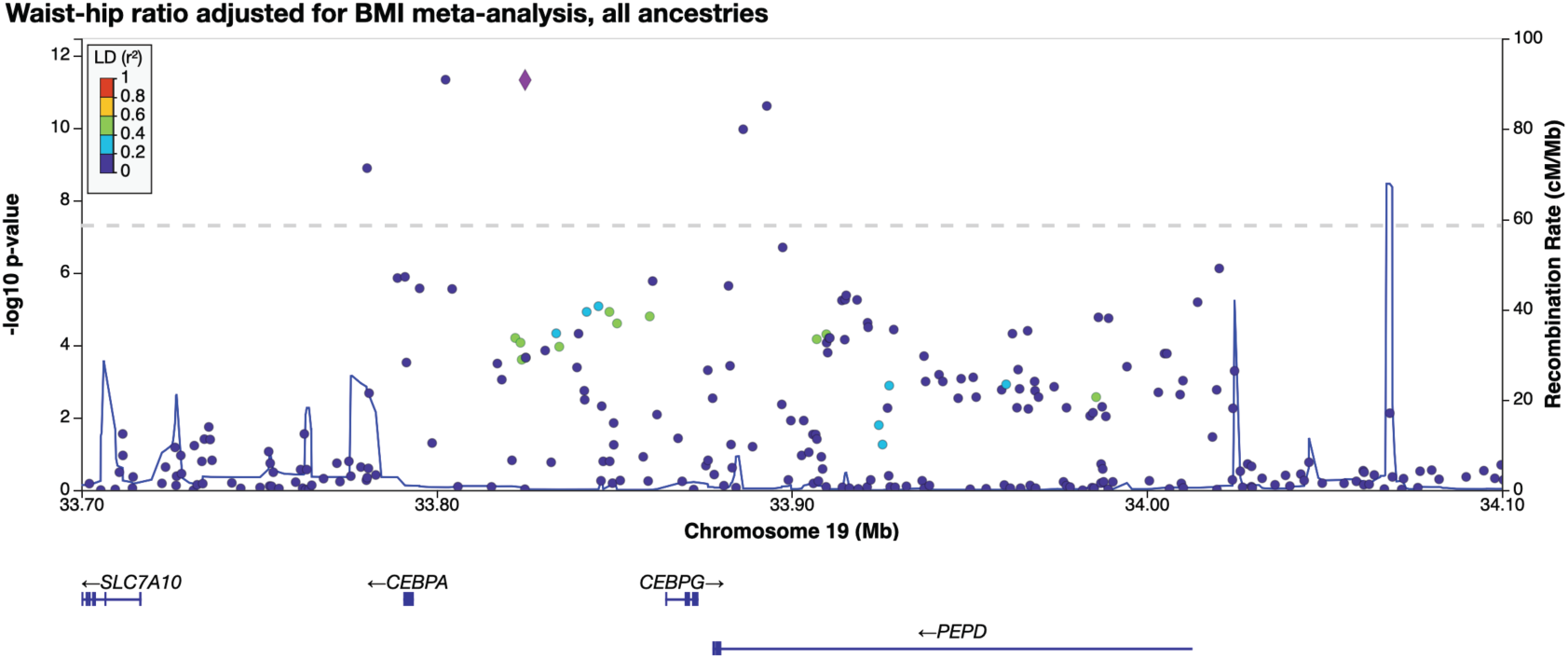
The *CEBPA* GWAS locus. LocusZoom plot of SNPs at the 19q13 locus associated with waist-to-hip ratio adjusted for BMI from the GIANT consortium [55].

**Supplementary Figure 2:**
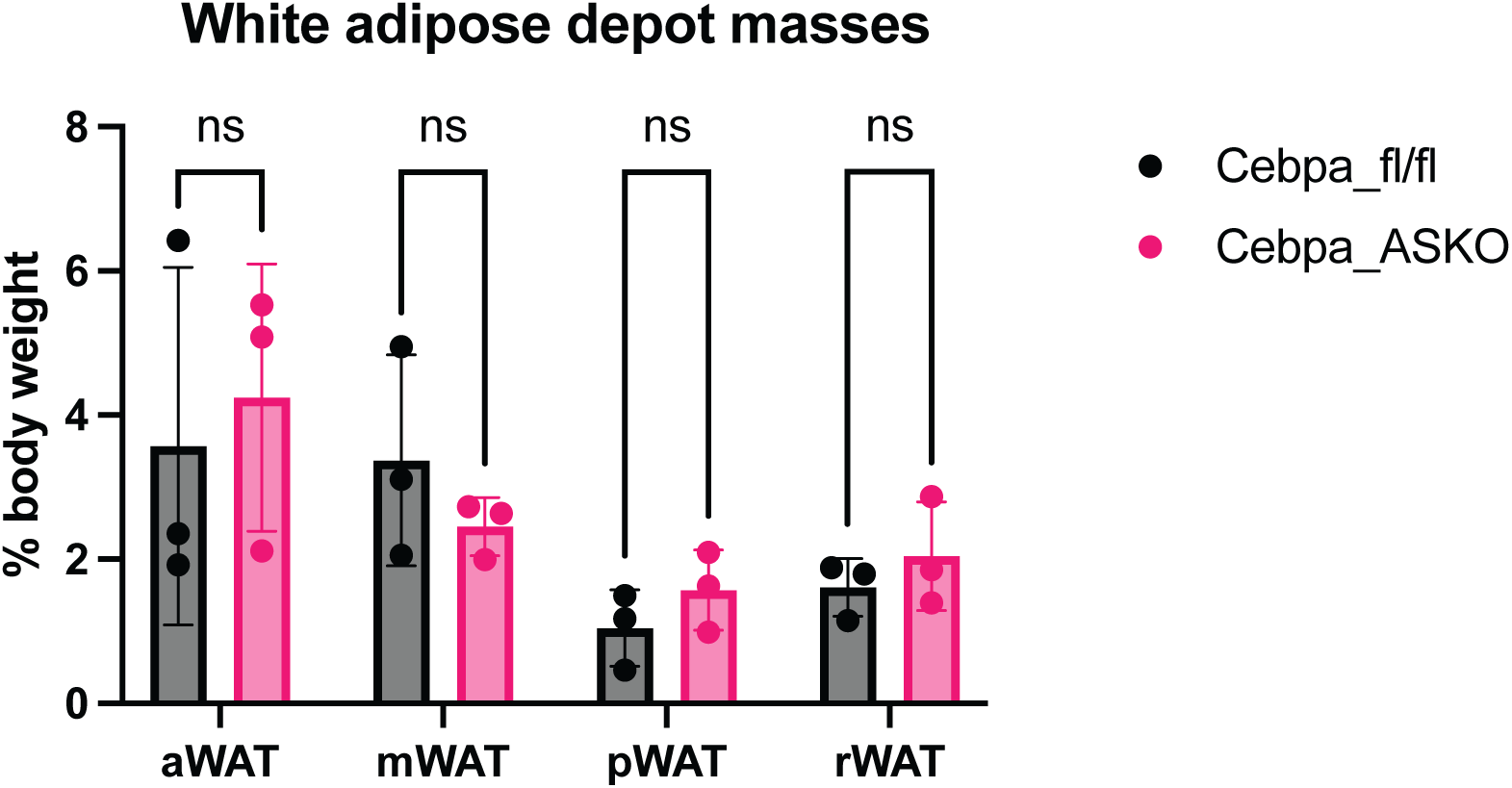
Cebpa_ASKO mice have no change in other WAT depot masses. **A.** Axillary, mesenteric, perirenal, and retroperitoneal adipose depot masses as percent body weight in male mice at 10-12 weeks of age (n=3). All mice were chow-fed. Student’s t-test was used to analyze results.

**Supplementary Figure 3:**
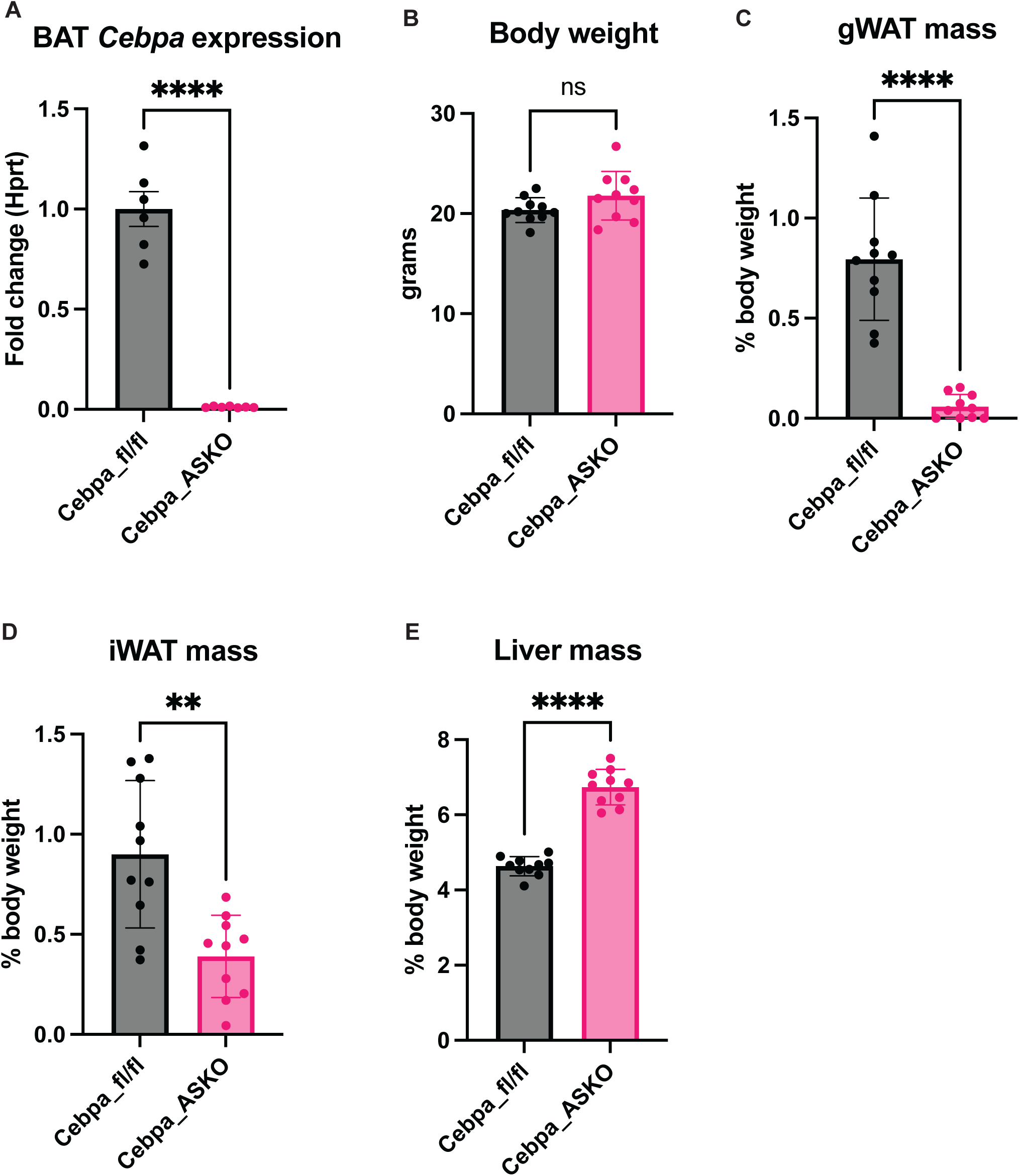
Female Cebpa_ASKO mice have reduced gWAT mass. **A.** TaqMan qPCR of *Cebpa* expression in (iwat) and BAT from 10-12-week-old female Cebpa_fl/fl and Cebpa_ASKO mice (n=6-7). **B.** Body weights at 10-12 weeks of female Cebpa_fl/fl and Cebpa_ASKO mice (n=10). **C-E.** gWAT, iWAT, and liver mass as percent body weight at 10-12 weeks of age (n=10). All mice were chow-fed. Student’s t-test was used to analyze results (**p<0.01, ****p<0.0001).

**Supplementary Figure 4:**
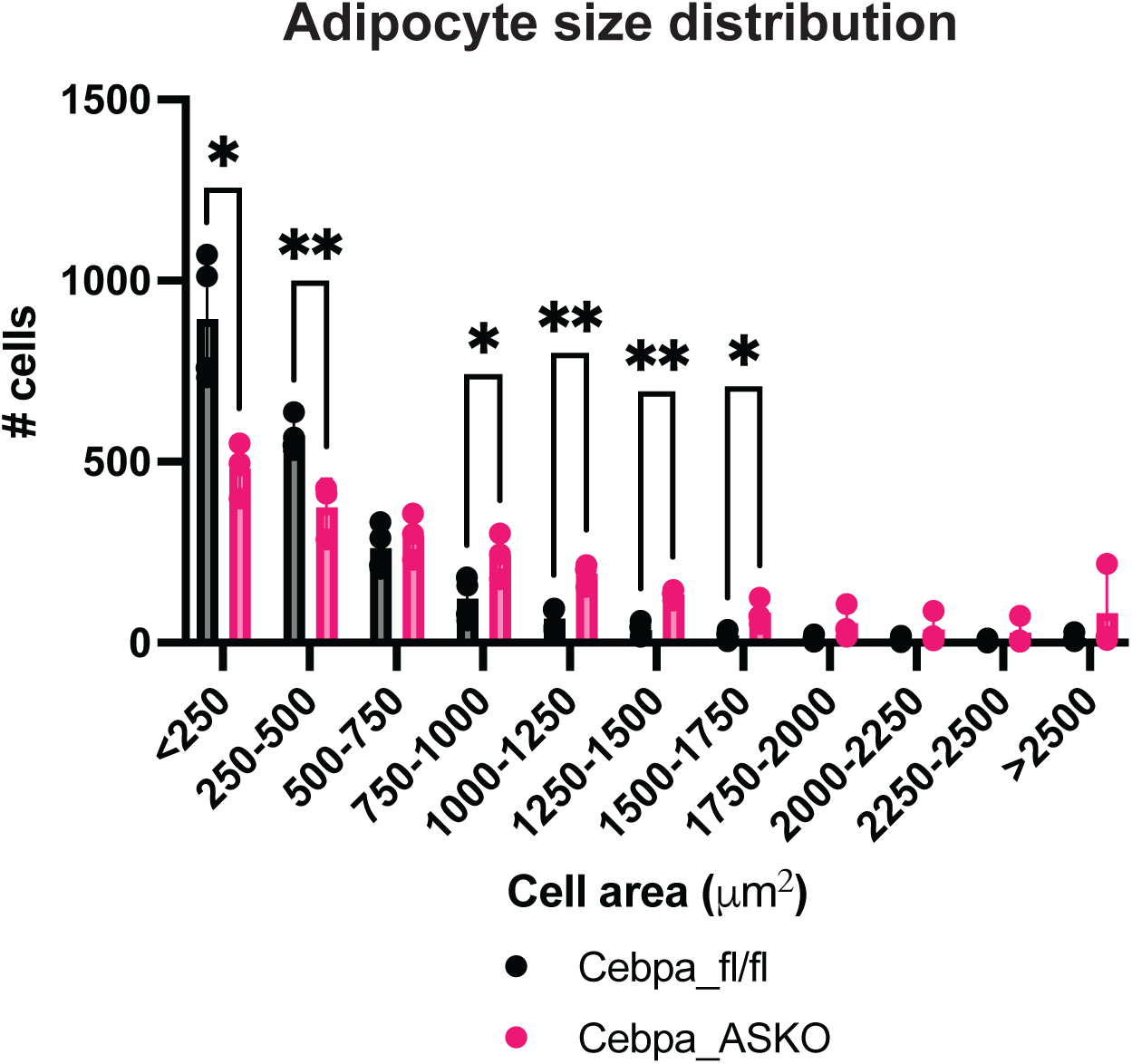
Cebpa_ASKO mice have larger iWAT adipocytes. **A**. Size distribution of adipocytes calculated using the Adiposoft FIJI plugin from representative iWAT H&E images of 10–12-week-old mice (n=3). All mice were chow-fed. Student’s t-test was used to analyze results (*p<0.05, **p<0.01).

**Supplementary Figure 5:**
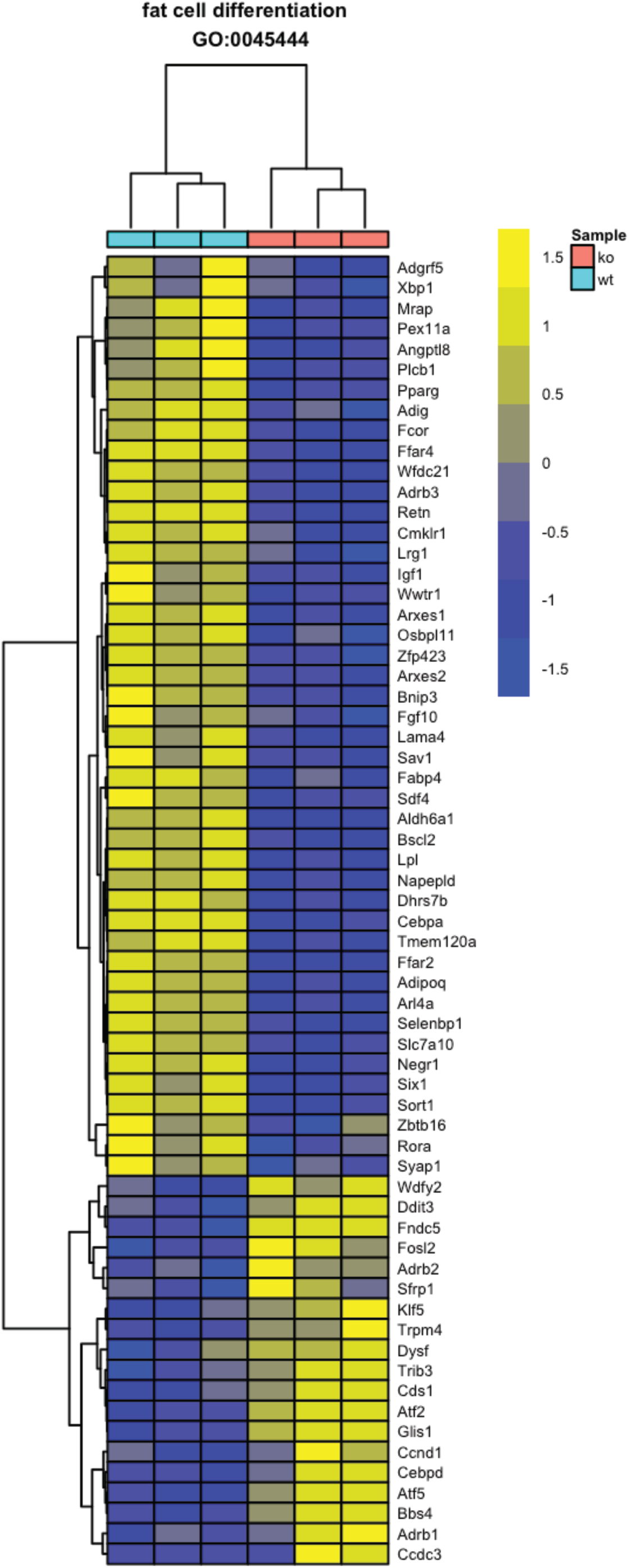
Cebpa_ASKO iWAT has changes in expression of adipocyte differentiation genes. Heatmap of expression of genes clustered in the fat cell differentiation GO term from bulk RNA-seq data (n=3). All mice were chow-fed.

**Supplementary Figure 6:**
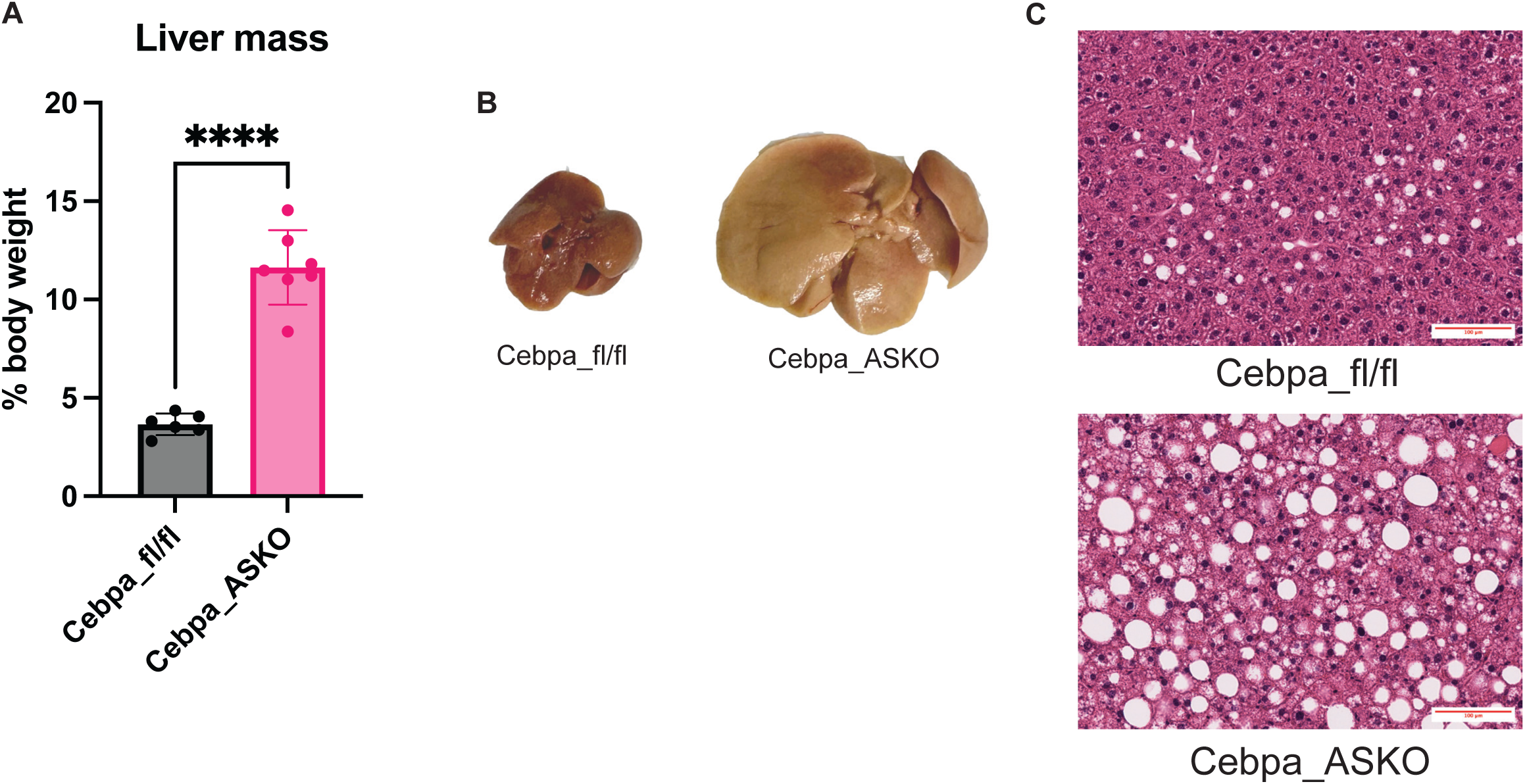
High fat diet feeding exacerbates the lipid deposition in livers of Cebpa_ASKO mice. **A**. Liver mass measured in HFD-fed Cebpa_fl/fl and Cebpa_ASKO mice. **B.** Macroscopic image of livers from HFD-fed Cebpa_fl/fl and Cebpa_ASKO mice. **C.** Representative images from H&E-stained livers. All mice were fed with HFD for 20 weeks. Student’s t-test was used to analyze results (****p<0.0001).

## Data availability statement

RNA-seq data will be available prior to publication from GEO (GSE302944). Other data that support the findings from this study are available from the corresponding author upon reasonable request.

## Bibliography

[1] Powell-Wiley, T.M., Poirier, P., Burke, L.E., Despres, J.P., Gordon-Larsen, P., Lavie, C.J., et al., 2021. Obesity and Cardiovascular Disease: A Scientific Statement From the American Heart Association. Circulation 143(21): e984–1010, Doi: 10.1161/CIR.0000000000000973.

[2] World Health Organization., 2024. Obesity and overweight.

[3] Neeland, I.J., Ayers, C.R., Rohatgi, A.K., Turer, A.T., Berry, J.D., Das, S.R., et al., 2013. Associations of visceral and abdominal subcutaneous adipose tissue with markers of cardiac and metabolic risk in obese adults. Obesity (Silver Spring) 21(9): E439–47, Doi: 10.1002/oby.20135.

[4] Rimm, E.B., Stampfer, M.J., Giovannucci, E., Ascherio, A., Spiegelman, D., Colditz, G.A., et al., 1995. Body Size and Fat Distribution as Predictors of Coronary Heart Disease among Middle-aged and Older US Men. American Journal of Epidemiology 141(12): 1117–27, Doi: 10.1093/oxfordjournals.aje.a117385.

[5] Ibrahim, M.M., 2010. Subcutaneous and visceral adipose tissue: structural and functional differences. Obes Rev 11(1): 11–8, Doi: 10.1111/j.1467-789X.2009.00623.x.

[6] McLaughlin, T., Lamendola, C., Liu, A., Abbasi, F., 2011. Preferential fat deposition in subcutaneous versus visceral depots is associated with insulin sensitivity. J Clin Endocrinol Metab 96(11): E1756–60, Doi: 10.1210/jc.2011-0615.

[7] Ruiz-Castell, M., Samouda, H., Bocquet, V., Fagherazzi, G., Stranges, S., Huiart, L., 2021. Estimated visceral adiposity is associated with risk of cardiometabolic conditions in a population based study. Scientific Reports 11(1): 9121, Doi: 10.1038/s41598-021-88587-9.

[8] Fu, J., Hofker, M., Wijmenga, C., 2015. Apple or pear: size and shape matter. Cell Metab 21(4): 507–8, Doi: 10.1016/j.cmet.2015.03.016.

[9] Loos, R.J.F., Yeo, G.S.H., 2022. The genetics of obesity: from discovery to biology. Nature Reviews Genetics 23(2): 120–33, Doi: 10.1038/s41576-021-00414-z.

[10] Shungin, D., Winkler, T.W., Croteau-Chonka, D.C., Ferreira, T., Locke, A.E., Magi, R., et al., 2015. New genetic loci link adipose and insulin biology to body fat distribution. Nature 518(7538): 187–96, Doi: 10.1038/nature14132.

[11] Speliotes, E.K., Willer, C.J., Berndt, S.I., Monda, K.L., Thorleifsson, G., Jackson, A.U., et al., 2010. Association analyses of 249,796 individuals reveal 18 new loci associated with body mass index. Nat Genet 42(11): 937–48, Doi: 10.1038/ng.686.

[12] Rosen, E.D., Walkey, C.J., Puigserver, P., Spiegelman, B.M., 2000. Transcriptional regulation of adipogenesis. Genes Dev 14(11): 1293–307.

[13] Cristancho, A.G., Lazar, M.A., 2011. Forming functional fat: a growing understanding of adipocyte differentiation. Nat Rev Mol Cell Biol 12(11): 722–34, Doi: 10.1038/nrm3198.

[14] Lefterova, M.I., Zhang, Y., Steger, D.J., Schupp, M., Schug, J., Cristancho, A., et al., 2008. PPARgamma and C/EBP factors orchestrate adipocyte biology via adjacent binding on a genome-wide scale. Genes Dev 22(21): 2941–52, Doi: 10.1101/gad.1709008.

[15] Linhart, H.G., Ishimura-Oka, K., DeMayo, F., Kibe, T., Repka, D., Poindexter, B., et al., 2001. C/EBPalpha is required for differentiation of white, but not brown, adipose tissue. Proc Natl Acad Sci U S A 98(22): 12532–7, Doi: 10.1073/pnas.211416898.

[16] Wang, Q.A., Tao, C., Gupta, R.K., Scherer, P.E., 2013. Tracking adipogenesis during white adipose tissue development, expansion and regeneration. Nat Med 19(10): 1338–44, Doi: 10.1038/nm.3324.

[17] Han, J., Lee, J.E., Jin, J., Lim, J.S., Oh, N., Kim, K., et al., 2011. The spatiotemporal development of adipose tissue. Development 138(22): 5027–37, Doi: 10.1242/dev.067686.

[18] Fang, H., Judd, R.L., 2018. Adiponectin Regulation and Function. Compr Physiol 8(3): 1031–63, Doi: 10.1002/cphy.c170046.

[19] Park, S., Oh, S.-Y., Lee, M.-Y., Yoon, S., Kim, K.-S., Kim, J., 2004. CCAAT/Enhancer Binding Protein and Nuclear Factor-Y Regulate Adiponectin Gene Expression in Adipose Tissue. Diabetes 53(11): 2757–66, Doi: 10.2337/diabetes.53.11.2757.

[20] Qiao, L., MacLean, P.S., Schaack, J., Orlicky, D.J., Darimont, C., Pagliassotti, M., et al., 2005. C/EBPα Regulates Human Adiponectin Gene Transcription Through an Intronic Enhancer. Diabetes 54(6): 1744–54, Doi: 10.2337/diabetes.54.6.1744.

[21] Straub, L.G., Scherer, P.E., 2019. Metabolic Messengers: Adiponectin. Nat Metab 1(3): 334–9, Doi: 10.1038/s42255-019-0041-z.

[22] Clarke, S.L., Robinson, C.E., Gimble, J.M., 1997. CAAT/enhancer binding proteins directly modulate transcription from the peroxisome proliferator-activated receptor gamma 2 promoter. Biochem Biophys Res Commun 240(1): 99–103, Doi: 10.1006/bbrc.1997.7627.

[23] Elberg, G., Gimble, J.M., Tsai, S.Y., 2000. Modulation of the murine peroxisome proliferator-activated receptor gamma 2 promoter activity by CCAAT/enhancer-binding proteins. J Biol Chem 275(36): 27815–22, Doi: 10.1074/jbc.M003593200.

[24] Thompson, B.R., Lobo, S., Bernlohr, D.A., 2010. Fatty acid flux in adipocytes: the in’s and out’s of fat cell lipid trafficking. Mol Cell Endocrinol 318(1–2): 24–33, Doi: 10.1016/j.mce.2009.08.015.

[25] Grabner, G.F., Xie, H., Schweiger, M., Zechner, R., 2021. Lipolysis: cellular mechanisms for lipid mobilization from fat stores. Nat Metab 3(11): 1445–65, Doi: 10.1038/s42255-021-00493-6.

[26] Cinti, S., Frederich, R.C., Zingaretti, M.C., De Matteis, R., Flier, J.S., Lowell, B.B., 1997. Immunohistochemical Localization of Leptin and Uncoupling Protein in White and Brown Adipose Tissue*. Endocrinology 138(2): 797–804, Doi: 10.1210/endo.138.2.4908.

[27] Zadeh, E.S., Lungu, A.O., Cochran, E.K., Brown, R.J., Ghany, M.G., Heller, T., et al., 2013. The Liver Diseases of Lipodystrophy: The Long-term Effect of Leptin Treatment. Journal of Hepatology 59(1): 131–7, Doi: 10.1016/j.jhep.2013.02.007.

[28] Savage, D.B., 2009. Mouse models of inherited lipodystrophy. Disease Models & Mechanisms 2(11–12): 554–62, Doi: 10.1242/dmm.002907.

[29] Polyzos, S.A., Perakakis, N., Mantzoros, C.S., 2019. Fatty liver in lipodystrophy: A review with a focus on therapeutic perspectives of adiponectin and/or leptin replacement. Metabolism 96: 66–82, Doi: 10.1016/j.metabol.2019.05.001.

[30] Jones, J.G., 2016. Hepatic glucose and lipid metabolism. Diabetologia 59(6): 1098–103, Doi: 10.1007/s00125-016-3940-5.

[31] Goldstein, J.L., Brown, M.S., 2015. A Century of Cholesterol and Coronaries: From Plaques to Genes to Statins. Cell 161(1): 161–72, Doi: 10.1016/j.cell.2015.01.036.

[32] Roshandel, D., Lu, T., Paterson, A.D., Dash, S., 2023. Beyond apples and pears: sex-specific genetics of body fat percentage. Front Endocrinol (Lausanne) 14: 1274791, Doi: 10.3389/fendo.2023.1274791.

[33] Hoffmann, T.J., Theusch, E., Haldar, T., Ranatunga, D.K., Jorgenson, E., Medina, M.W., et al., 2018. A large electronic-health-record-based genome-wide study of serum lipids. Nature Genetics 50(3): 401–13, Doi: 10.1038/s41588-018-0064-5.

[34] Spracklen, C.N., Iyengar, A.K., Vadlamudi, S., Raulerson, C.K., Jackson, A.U., Brotman, S.M., et al., 2020. Adiponectin GWAS loci harboring extensive allelic heterogeneity exhibit distinct molecular consequences. PLoS Genetics 16(9): e1009019, Doi: 10.1371/journal.pgen.1009019.

[35] Barton, A.R., Sherman, M.A., Mukamel, R.E., Loh, P.R., 2021. Whole-exome imputation within UK Biobank powers rare coding variant association and fine-mapping analyses. Nat Genet 53(8): 1260–9, Doi: 10.1038/s41588-021-00892-1.

[36] Hubel, C., Gaspar, H.A., Coleman, J.R.I., Finucane, H., Purves, K.L., Hanscombe, K.B., et al., 2019. Genomics of body fat percentage may contribute to sex bias in anorexia nervosa. Am J Med Genet B Neuropsychiatr Genet 180(6): 428–38, Doi: 10.1002/ajmg.b.32709.

[37] Martin, S., Cule, M., Basty, N., Tyrrell, J., Beaumont, R.N., Wood, A.R., et al., 2021. Genetic Evidence for Different Adiposity Phenotypes and Their Opposing Influences on Ectopic Fat and Risk of Cardiometabolic Disease. Diabetes 70(8): 1843–56, Doi: 10.2337/db21-0129.

[38] Sakaue, S., Kanai, M., Tanigawa, Y., Karjalainen, J., Kurki, M., Koshiba, S., et al., 2021. A cross-population atlas of genetic associations for 220 human phenotypes. Nat Genet 53(10): 1415–24, Doi: 10.1038/s41588-021-00931-x.

[39] Liu, Y., Basty, N., Whitcher, B., Bell, J.D., Sorokin, E.P., van Bruggen, N., et al., 2021. Genetic architecture of 11 organ traits derived from abdominal MRI using deep learning. Elife 10, Doi: 10.7554/eLife.65554.

[40] Graff, M., Scott, R.A., Justice, A.E., Young, K.L., Feitosa, M.F., Barata, L., et al., 2017. Genome-wide physical activity interactions in adiposity - A meta-analysis of 200,452 adults. PLoS Genet 13(4): e1006528, Doi: 10.1371/journal.pgen.1006528.

[41] Pulit, S.L., Stoneman, C., Morris, A.P., Wood, A.R., Glastonbury, C.A., Tyrrell, J., et al., 2019. Meta-analysis of genome-wide association studies for body fat distribution in 694 649 individuals of European ancestry. Hum Mol Genet 28(1): 166–74, Doi: 10.1093/hmg/ddy327.

[42] Zhu, Z., Guo, Y., Shi, H., Liu, C.L., Panganiban, R.A., Chung, W., et al., 2020. Shared genetic and experimental links between obesity-related traits and asthma subtypes in UK Biobank. J Allergy Clin Immunol 145(2): 537–49, Doi: 10.1016/j.jaci.2019.09.035.

[43] Pellegrinelli, V., Rodriguez-Cuenca, S., Rouault, C., Figueroa-Juarez, E., Schilbert, H., Virtue, S., et al., 2022. Dysregulation of macrophage PEPD in obesity determines adipose tissue fibro-inflammation and insulin resistance. Nat Metab 4(4): 476–94, Doi: 10.1038/s42255-022-00561-5.

[44] Wang, Q.A., Tao, C., Jiang, L., Shao, M., Ye, R., Zhu, Y., et al., 2015. Distinct regulatory mechanisms governing embryonic versus adult adipocyte maturation. Nat Cell Biol 17(9): 1099–111, Doi: 10.1038/ncb3217.

[45] Christy, R.J., Yang, V.W., Ntambi, J.M., Geiman, D.E., Landschulz, W.H., Friedman, A.D., et al., 1989. Differentiation-induced gene expression in 3T3-L1 preadipocytes: CCAAT/enhancer binding protein interacts with and activates the promoters of two adipocyte-specific genes. Genes Dev 3(9): 1323–35, Doi: 10.1101/gad.3.9.1323.

[46] Ambele, M.A., Dhanraj, P., Giles, R., Pepper, M.S., 2020. Adipogenesis: A Complex Interplay of Multiple Molecular Determinants and Pathways. Int J Mol Sci 21(12), Doi: 10.3390/ijms21124283.

[47] Rondini, E.A., Granneman, J.G., 2020. Single cell approaches to address adipose tissue stromal cell heterogeneity. Biochem J 477(3): 583–600, Doi: 10.1042/BCJ20190467.

[48] Nahmgoong, H., Jeon, Y.G., Park, E.S., Choi, Y.H., Han, S.M., Park, J., et al., 2022. Distinct properties of adipose stem cell subpopulations determine fat depot-specific characteristics. Cell Metab 34(3): 458–472 e6, Doi: 10.1016/j.cmet.2021.11.014.

[49] Sanchez-Gurmaches, J., Hung, C.M., Guertin, D.A., 2016. Emerging Complexities in Adipocyte Origins and Identity. Trends Cell Biol 26(5): 313–26, Doi: 10.1016/j.tcb.2016.01.004.

[50] Hong, K.Y., Bae, H., Park, I., Park, D.Y., Kim, K.H., Kubota, Y., et al., 2015. Perilipin+ embryonic preadipocytes actively proliferate along growing vasculatures for adipose expansion. Development 142(15): 2623–32, Doi: 10.1242/dev.125336.

[51] Maniyadath, B., Zhang, Q., Gupta, R.K., Mandrup, S., 2023. Adipose tissue at single-cell resolution. Cell Metab 35(3): 386–413, Doi: 10.1016/j.cmet.2023.02.002.

[52] Zhang, Q., Shan, B., Guo, L., Shao, M., Vishvanath, L., Elmquist, G., et al., 2022. Distinct functional properties of murine perinatal and adult adipose progenitor subpopulations. Nat Metab 4(8): 1055–70, Doi: 10.1038/s42255-022-00613-w.

[53] Sarvari, A.K., Van Hauwaert, E.L., Markussen, L.K., Gammelmark, E., Marcher, A.B., Ebbesen, M.F., et al., 2021. Plasticity of Epididymal Adipose Tissue in Response to Diet-Induced Obesity at Single-Nucleus Resolution. Cell Metab 33(2): 437–453 e5, Doi: 10.1016/j.cmet.2020.12.004.

[54] Hepler, C., Shan, B., Zhang, Q., Henry, G.H., Shao, M., Vishvanath, L., et al., 2018. Identification of functionally distinct fibro-inflammatory and adipogenic stromal subpopulations in visceral adipose tissue of adult mice. Elife 7, Doi: 10.7554/eLife.39636.

[55] Pruim, R.J., Welch, R.P., Sanna, S., Teslovich, T.M., Chines, P.S., Gliedt, T.P., et al., 2010. LocusZoom: regional visualization of genome-wide association scan results. Bioinformatics 26(18): 2336–7, Doi: 10.1093/bioinformatics/btq419.

